# Stress Resistance Heat Shock Protein 70 (HSP70) Analysis in Sorghum (*Sorghum bicolor* L.) at Genome-Wide Level

**DOI:** 10.1101/2021.11.22.469496

**Authors:** Kasahun Amare, Mulugeta Kebede

**Affiliations:** Department of Plant Science, College of Agriculture and Natural Resource, Mekdela Amba University, Tuluawlia, Ethiopia; Department of Applied Biology, School of Applied Natural Science, Adama Science and Technology University, Adama, Ethiopia

**Keywords:** Gene structure, Cis-acting elements, Promoter region, Protein motif

## Abstract

Heat shock proteins (HSP70) play an important role in many biological processes. However, as typical in Sorghum bicolor, the systematic identification of the HSP70 gene is very limited, and the role of the Hsp70 gene in the evolution of Sorghum bicolor has not been described systematically a lot. To overcome the gap, Insilco analysis of HSP70 gene family was conducted.The investigation was utilizing the bioinformatics method to analyze the HSP70 gene family and it has been identified that 30 HSP70 genes from the genome sequence of Sorghum bicolor. A comprehensive analysis of these 30 identified genes undertaking the analysis of gene structure, phylogeny, and physicochemical properties, subcellular localization, and promoter region analysis. The gene structure visualization analyses revealed that 22 genes contains both 5’ and 3’ UTRS and one 5’ and one 3’ gene and 6 genes without UTR. The highest number of introns was recorded as 12 and those genes have shown that without in any intron. In the promoter region analysis, ten protein motifs are identified and characterized and 2219 cis-acting elements are identified. Among those, the promoter enhancer elements share the highest number (1411) and light-responsive elements share the next value (335). The physicochemical properties analysis revealed that 23 families have an acidic nature while four families are basic and the rests are neutral. In general, the different analyses performed disclosed their structural organization, subcellular localization, physicochemical properties, cis-acting elements, phylogenetic, and understress conditions. This study provides further information for the functional characterization of HSP70 and helps to understand the mechanisms of abiotic stress tolerance under diverse stress conditions in Sorghum bicolor.

## 1. INTRODUCTION

Sorghum (*Sorghum bicolor* L.) is a multipurpose food crop belonging to the Poaceae family, which are C4 carbon cycle plants with high photosynthetic efficiency and productivity. It ranks the top five among cereal crops in the world. It serves as a source of food, fodder, feed, and bioenergy [2]. It plays a major function in the food security of sub-Saharan Africa, supporting about 500 million citizens. It is cultivated in drought-prone and subsidiary areas in semi-arid zones where other crops cannot grow constantly [1, 17].

The most critical abiotic stresses that hamper the production of sorghum harvests may comprise: supplement inadequacy, aluminum stress, dry season, high saltiness, waterlogging, and temperature stress, these wonders must be adapted to plants during development [39].

The effects of stress can prompt inadequacies in development, crop yields, perpetual harm, or passing if the stress surpasses the plant resilience limits [25]. The endurance of plants is restricted to an expected warm scope of −10 to +60°C, characterized by the edge of freezing over intracellular water and the temperature of protein denaturation [38]. Such abiotic stresses have been uncovered to cause the development of different intracellular substances, including nucleic acids, amino acids, starches, and proteins. Following the kickoff of molecular biology strategies into plant science, an enormous arrangement of exertion went into the disclosure of stress-inducible genes [13].

Along these lines, an understanding of the abiotic stress response is presently thought to be quite possibly the main point in plant science. The main headway in this exploration field has come from the use of molecular biology techniques. After this method was utilized in plant science, different abiotic stress-inducible genes were isolated and their function was correctly characterized in transgenic plants. The accessibility of this information widened and extended our perspective on abiotic stress response and resilience in plants [13]. In such a manner, plants react to abiotic stress like heat also as different anxieties that can trigger plant genes to protect the strange conditions of gene articulation that was not expressed under ordinary conditions. Such sort of reaction with weights on the molecular level is likewise found in living frameworks enveloping microorganisms, plants, and animals [12, 35].

A group of genes incited during heat stress and expressed proteins are called heat-shock proteins (HSPs), stress-prompting proteins, or stress proteins [10]. HSPs are known to be expressed in plants when they experience high-temperature stress as well as in light of a wide scope of other ecological burdens, for example, water stress, salt stress, cold stress, and oxidative stress [39, 43].

These stress reacting proteins are classified into various groups dependent on their function and expression pattern, such as constitutive heat stun proteins that are expressed constitutively, and inducible structures that are expressed in light of specific components [4]. Besides, it has been grouped dependent on their protein molecular weight, where they are partitioned into HSP90 (83~110 kDa), HSP70 (66~78 kDa), HSP60 (58~65 kDa), and other small molecular weight proteins HSP20s [44]. HSP70 is described by two utilitarian domains, the amino N-terminal ATPase domain (44 kDa) showing an ATPase activity and a carboxyl C-terminal peptide-restricted domain (25 kDa). The peptide-restricting domain is additionally partitioned into a β-sandwich subdomain (18 kDa), which is the substrate-restricting domain and an α-helical subdomain [26, 47]. The 70-kDa heat shock proteins (HSP70s) are the most bountiful and generally considered a conserved group of proteins. sHSPs comprise small molecular weight proteins that function as molecular chaperones basic for protein collapsing and avoidance of irreversible protein aggregation [44]. Notwithstanding sHSPs being unmistakably expressed during the heat stun reactions in plants, it is presently realized that some are expressed in unstressed cells also and are thusly engaged with measures other than heat stress [40; 44]. For instance, in plants, they are upregulated during the ripening initiation of tomato fruit [37] and may likewise secure ready tomato fruit against chilling injury [31]. The small utilitarian domains sHsps/Hsp20s have been distinguished in sorghum as molecular chaperones that keep up appropriate folding, trafficking, and disaggregation of proteins under assorted abiotic stress conditions [21].

### 1.1. The Problem Statement

The HSPs being prominently expressed during the heat shock response in plants is now known that some are expressed in unstressed cells as well and are, therefore, involved in processes other than heat stress [40; 44]. Several studies have revealed that HSP70 is closely associated with plant abiotic stress [41], disease resistance [22], growth, and development [42]. Heat tolerance in various plant species such as *Triticum aestivum*[7], *Oryza sativa* [35], and *Capsicum annuum* [9]. Small molecular weight HSPs function as molecular chaperones critical for protein folding and prevention of irreversible protein aggregation [44]. When the plant suffers from high temperature, drought, high salt, low temperature, and heavy metals, HSP70s rapidly accumulate to maintain the stability of the protein and biological macromolecules to improve the resistance of the plant [43]. In addition, some studies found that the HSP protein has some relationship with plant embryogenesis. They are also upregulated during the ripening initiation of tomato fruit [37] and may also protect ripe tomato fruits against chilling injury [31]. The relationship between heat shock treatment and embryogenesis was also studied in *Brassica napus*, and HSP70 and HSP90 located in the nucleus and cytoplasm were found to be rapidly induced [36].

Despite the breakthrough in the identification and characterization of plant HSPs in sorghum, the most widely cultivated and stress-tolerant cereal crop in the world and particularly sub-Saharan countries [17], the mechanism of stress tolerance through HSP70 as quite a few studies have been reported before. Thus, the identification and characterization of HSP70 genes in sorghum will aid in a better understanding of the molecular mechanism of its stress tolerance. In sorghum, the existence of HSP70 was verified by immunoblotting following salt stress [28–29], and heat stress [26], and Nagaraju et al. [27] identified sHSP family which uses as chaperons. However, the authors could’t find any literature describing the HSP70 gene characterization at the genomewide level. Due to the effect of various stresses on plants, investigating various mechanisms of plant stress responses is crucial. Thus, a genome-wide level analysis of sorghum HSP70 gene will help to reveal the underlying complex molecular mechanisms. The publication of the genome data of sorghum will enable systematic analyses of HSP70 evolution and function.

In this study, the bioinformatics method was used to analyze genomic HSP70 gene family members of sorghum, including the number of gene identification, phylogenetic relationships, gene structural features (exon-intron organization), and subcellular localization of the HSP70 protein.

### 1.2. Objectives of the Study

#### 1.2.1. General Objective

- To perform stress resistance gene family analysis in sorghum at a genome-wide level

#### 1.2.2 Specific Objectives

- To identify the HSP70 genes from the database
- To perform phylogenetic relationship analysis
- To perform gene structural visualization
- To perform the physicochemical properties of the HSP70 protein family
- To perform HSP70 gene motif and cis-acting element analysis

### 1.3. Significance of the Study

This study’s findings will redound to society’s benefit, considering that plant breeders play a vital role in crop science and plant biotechnology /crop improvement/ today. The greater demand for graduates with plant genetics, plant breeding, and plant biotechnology backgrounds justifies the need for more effective, life-changing technology development. Thus, organizations that apply the recommended approach derived from the results of this study will benefit better. Because one of the breeding methods that tries to find a solution for various breeding bottlenecks are the molecular approaches. As a result, understanding sorghum HSP70s has great importance since sorghum is severely affected by heat stress, particularly during the grain filling stage. Therefore, after the completion of the groundwork, it would be useful for the functional identification of the HSP70 gene and its application in breeding more adaptable sorghum cultivars’ development and engineering of the genes.

## 2. MATERIALS AND METHODS

### 2.1. Study Materials

The study materials in this research have consisted of the proteomic and genomic sequences of sorghum, maize, wheat, rice, sugarcane, millet, and Arabidopsis. Others like online and offline based software including, TBtool, MEGAX, Microsoft excel, and web based databases like NCBI, MEME, PlantCARE, Pfam, GSDS, Phytozome, ExPasy, and Cello life were used.

### 2.2. Database Mining (HSP70 Family Genes)

The whole sorghum genome sequence was downloaded from the annotation database phytozome V3.1.1 (Cereal grass) database using a query sequence that was obtained from NCBI(Table1). To gather the probable candidates’ sorghum HSP70 protein sequence, the HMM profile of the HSP70 domain was first checked in the Pfam database. The sorghum gene sequence information, including; the gene coordinate in the chromosome, genome sequence, full CDS sequence, protein sequence, and 2k bp of the nucleotide sequences upstream of the translation initiation codon was downloaded from the phytozome. The molecular weight (kDa) and isoelectric point (PI), the grand average hydropathy (GRAVY) value of each gene was calculated using the compute PI/MW tool from EXPASY proteome server (https://web.expasy.org/cgi-bin/protparam/protparam). Consequently, the genes were renamed as SbSHP70-1 to SbSHP70-30 for convenience, and different parameters were analyzed as depicted in Table-2.

**Table 1:** HSP70 Gene Family Sequences Identified From Phytozome and NCBI.

| From phytozome( with accession number) |  |  |  | From NCBI(with accession number) |
| --- | --- | --- | --- | --- |
| *Sorghum bicolor* |  | *Arabidopsis thaliana* |  | *Others* |
| Sobic.001G118600 | Sobic.002G249800 | AT1G09080 | T5G28540 | *Triticum aestivum* (AHW48175.1) |
| Sobic.001G129000 | Sobic.003G178101 | AT1G11660 | T5G42020 | *Zea mays* (ACG29713.1) |
| Sobic.001G193500 | Sobic.003G350700 | AT1G16030 | T5G49910 | *Panicum miliaceum* (RLN16627.1) |
| Sobic.001G418600 | Sobic.003G378700 | AT1G56410 |  | *Saccharum officinarum* (ANN87873.1) |
| Sobic.001G419000 | Sobic.004G011700 | AT1G79920 |  | *Oryza sativa* (ABF93623.1) |
| Sobic.001G419300 | Sobic.004G263500 | AT1G79930 |  |  |
| Sobic.001G419400 | Sobic.006G055600 | AT2G32120 |  |  |
| Sobic.001G419500 | Sobic.008G038700 | AT3G09440 |  |  |
| Sobic.001G419600 | Sobic.008G088200 | AT3G12580 |  |  |
| Sobic.001G419700 | Sobic.008G136000 | AT4G16660 |  |  |
| Sobic.001G419801 | Sobic.009G066900 | AT4G24280 |  |  |
| Sobic.001G419900 | Sobic.009G067000 | AT4G37910 |  |  |
| Sobic.001G420100 | Sobic.009G163900 | AT5G02490 |  |  |
| Sobic.001G454000 | Sobic.009G254800 | AT5G02500 |  |  |
| Sobic.002G008000 | Sobic.010G230600 | AT5G09590 |  |  |

**Table 2:**
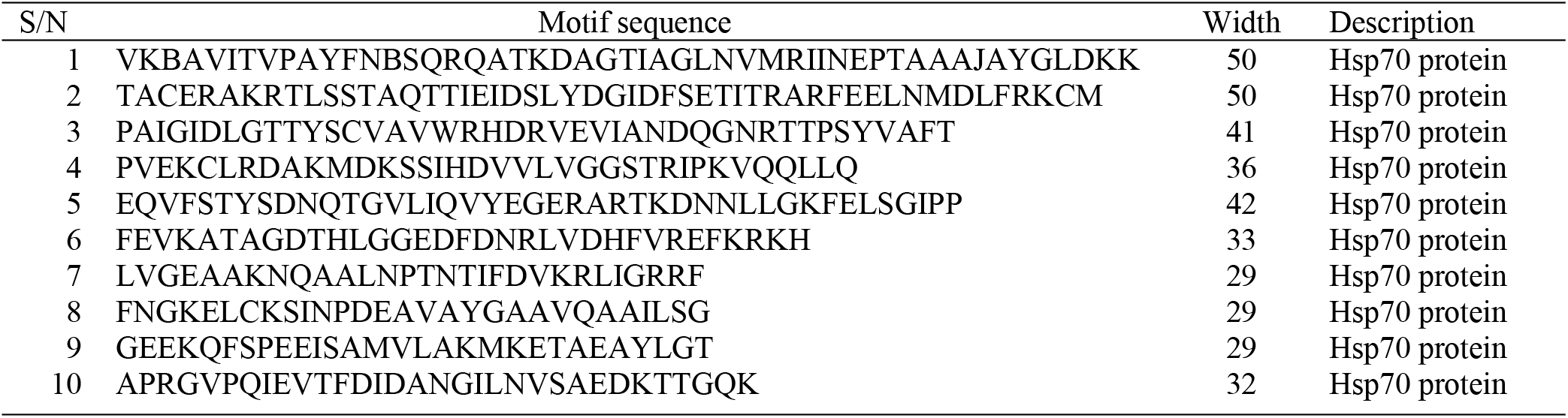
Conserved motif result as predicted by meme.

### 2.3. Gene Structure Display for HSP70 Genes

The Gene Structure Display Server tool (http://gsds.gao-lab.org/index.php) was used to analyze the exon-intron structures [14]. Besides the exon and intron regions, the upstream and downstream UTR (un-translated) regions were also determined to show possible structures of entirely expressed mRNA. Intron phases were classified based on their positions relative to the reading frame of the translated proteins: phase 0 (located between two codons), phase 1 (splitting codons between the first and second nucleotides), or phase 2 (splitting codons between the second and third nucleotides).

### 2.4. Protein Motif and Cis Regulatory Elements Prediction of HSP70 Genes

To find conserved motifs in sorghum Hsp70 gene family members, The MEME suite (version 5.3.3) (https://meme-suite.org/meme/tools/meme) [3] was used to search for motifs in all HSP70 genes that was downloaded from phytozome.

To investigate the cis-acting elements, the upstream regions of all HSP70 genes (2kbp) were extracted from the Phytozome website (https://phytozome.jgi.doe.gov/pz/portal.html). Subseque ntly, all of the sequences were submitted to the PlantCARE website (http://bioinformatics.psb.ugent.be/webtools/plantcare/html/) to identify possible cis-acting elements.

### 2.5. Phylogenetic Analysis

HSP70 protein sequences for phylogenetic analyses were collected from related plant species of sorghum. These species, including *Zea mays, Oryza sativa, and Triticum aestivum, Saccharum officinarum, Panicum miliaceum*, and the model plant *Arabidopsis thaliana*. Their HSP70 protein sequences were downloaded from the NCBI database and that of *Sorghum bicolor* HSP70 sequence was downloaded from Phytozome V3.1.1 (Cereal grass) database. The multiple sequences alignments of the HSP70 domains were done with ClustalW in MEGAX (version 10.05) with default settings and to construct phylogenetic trees Neighbor-Joining method with 1,000 bootstrap replicates was followed.

### 2.6. Gene Localization and Gene Duplication Analysis

The amino acid sequences of tandemly and segmentally duplicated HSP70 genes were subjected to TBtool for the analysis of synonymous (Ks) and nonsynonymous (Ka) substitution rate determination. In light of a pace of 6.1 x 10^-9^ replacements for each site each year, the difference time (T) was determined as T= Ks/(2 x 6.1 10^-9^) x 10^-6^ million years prior (Mya). The gene location on the chromosome was analyzed with the help of <u>Phenogram - Ritchie Lab</u> to annotate with lines in color at specific base pair locations. PhenoGram allows for annotation of chromosomal locations and/or regions with shapes in different colors, gene identifiers, or other text.

### 2.7. Protein Sub-Cellular Localization Prediction and Physico-Chemical Properties

Protein subcellular localization is crucial for genome annotation and protein function prediction. Therefore, the subcellular localization of proteins was analyzed using cello life. For computing, physicochemical features such as molecular mass, isoelectric point, instability index, aliphatic index, and average hydropathy were computed using ProtParam expasy (https://web.expasy.org/protparam).

## 3. RESULTS AND DISCUSSION

### 3.1. Identification of the HSP70 Gene Family in Sorghum bicolor

The systematic searching of the sorghum genome showed that a total of 39 genes were responsible for SbHSP70 protein production/ coding. However, among the identified genes, eight genes were found as redundant versions of other genes and one gene was a fragmented sequence and therefore was removed. After the removal of redundant sequences, domains in the proteins of these gene families were searched using the Pfam search tool for confirming the presence of specific Hsp70 domains. Under the activity, a total of nine genes were removed and 30 genes were identified. All nonredundant HSP70 genes were distributed on chromosomes 1, 2, 3, 4, 6, 8, 9, and 10 of sorghum. The largest numbers of genes (14) are distributed on chromosome one and four genes on chromosome nine (Table 3 and Figure 3). It was understood from different kinds of literature previously various numbers of HSP70 genes were identified in different plant species, for instance, 17 Hsp70 in *Hordeum vulgare* [5], 20 StHSP70 in *Solanum tuberosum* [19] and 21 CaHSP70 genes in *Capsicum annuum* [8], *24* PvHSP70 *in Phaseolus vulgaris* and 61 HSP70 in *Glycine max* were also reported [44]. This variation in HSP70s family in plants may be due to the presence of extra organelles, like plastids, in the plant cell compared to other eukaryotic organisms.

**Table 3:**
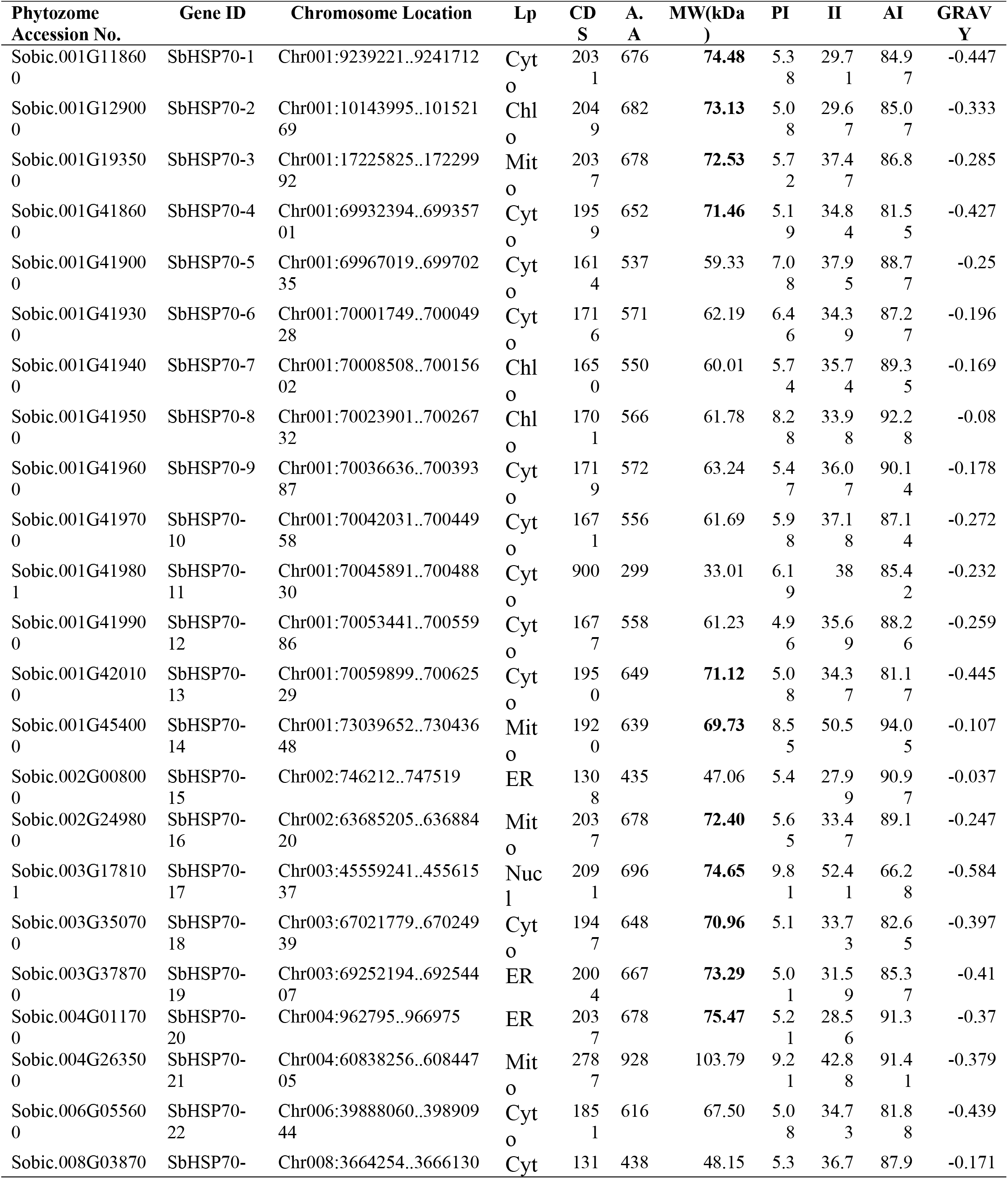
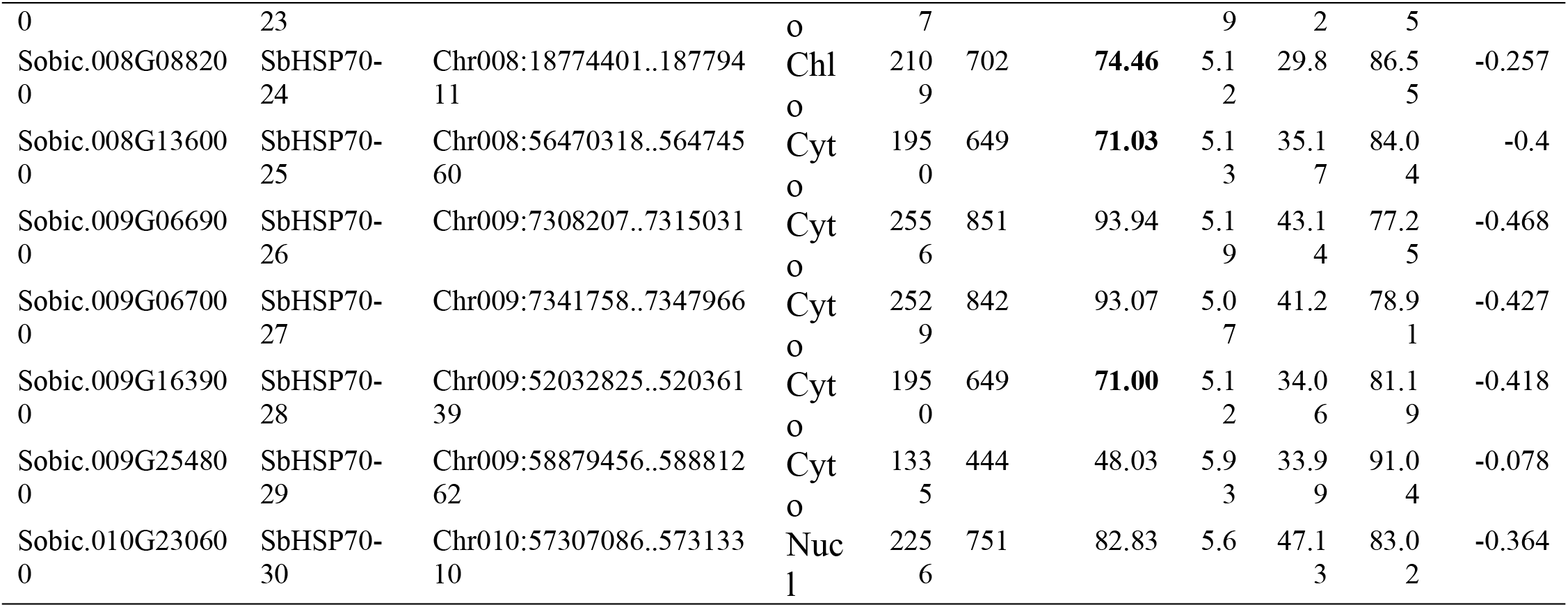
Physico-chemical properties and subcellular localization of HSP70 protein.

### 3.2. Gene Structure Display for HSP70 Genes

The gene structure was determined for 30 SbHSP70 gene families using both genomic sequences and coding sequences. From the analysis, it has been noticed that a total of 22 genes were found to have both UTRs (5’ and 3’ end) and two genes were found with only one UTR each(either 5’ or 3’), and six genes were found to have no UTR. The maximum number of introns were found in Sobic.004G263500 gene, that is, 12, followed by eight introns in Sobic.004G011700, Sobic.009G066900, Sobic.009G067000, and Sobic.010G230600 genes respectively. Furthermore, no introns were found in two SbHSP70s genes (Sobic.002G008000 and Sobic.003G378700) (Figure-1 and Table-4).

**Figure 1:**
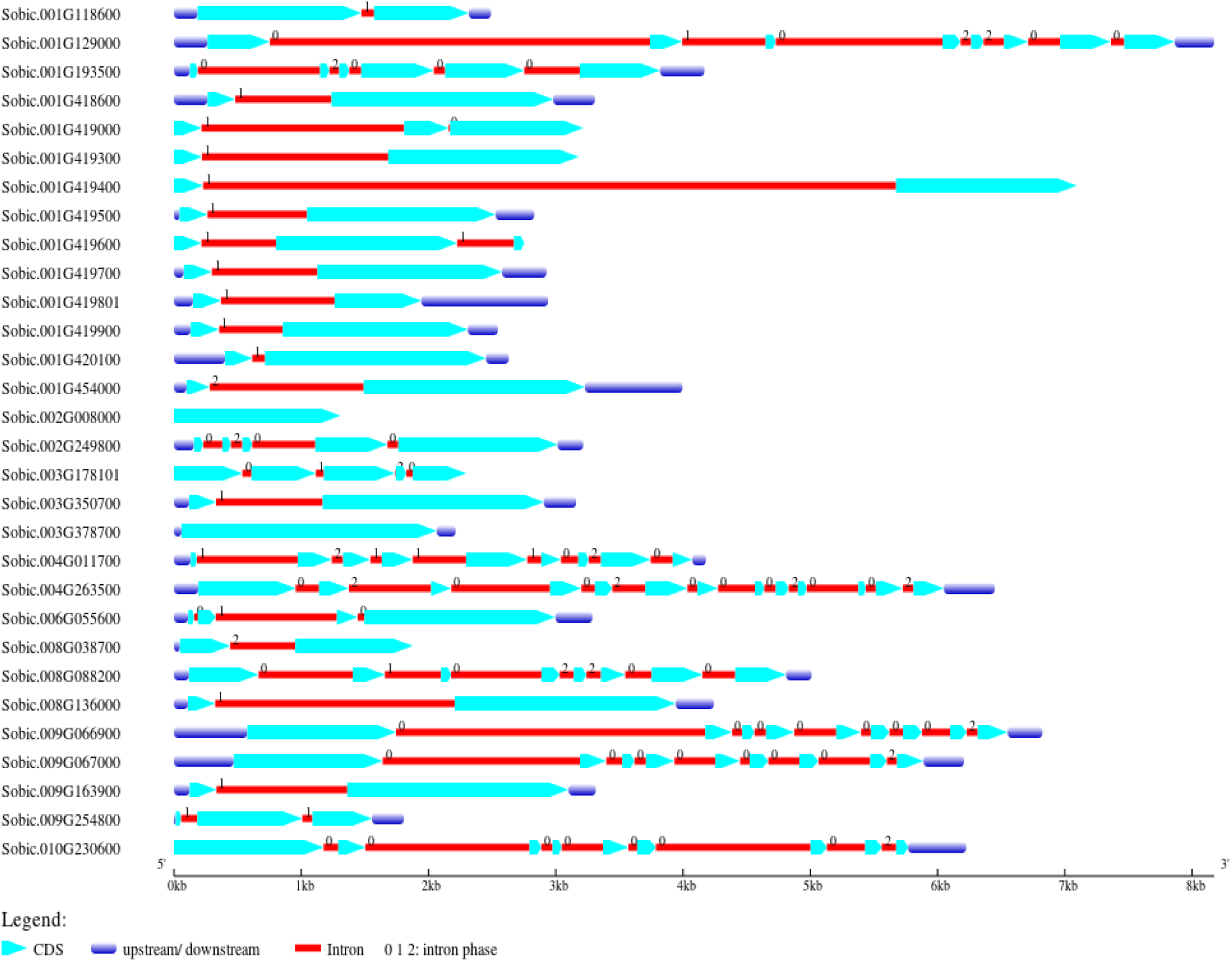
Exon/intron organization of 30 HSP70s genes in sorghum.

**Table 4:** Physico-chemical properties and subcellular localization of HSP70 protein (cont…)

| Phytozome Accession No. | Gene ID | Chromosome Location | Lp | Exon/Intron |
| --- | --- | --- | --- | --- |
| Sobic.001G118600 | SbHSP70-1 | Chr001:9239221..9241712 | Cyto | 2:01 |
| Sobic.001G129000 | SbHSP70-2 | Chr001:10143995..10152169 | Chlo | 8:07 |
| Sobic.001G193500 | SbHSP70-3 | Chr001:17225825..17229992 | Mito | 6:05 |
| Sobic.001G418600 | SbHSP70-4 | Chr001:69932394..69935701 | Cyto | 2:01 |
| Sobic.001G419000 | SbHSP70-5 | Chr001:69967019..69970235 | Cyto | 3:02 |
| Sobic.001G419300 | SbHSP70-6 | Chr001:70001749..70004928 | Cyto | 2:01 |
| Sobic.001G419400 | SbHSP70-7 | Chr001:70008508..70015602 | Chlo | 2:01 |
| Sobic.001G419500 | SbHSP70-8 | Chr001:70023901..70026732 | Chlo | 2:01 |
| Sobic.001G419600 | SbHSP70-9 | Chr001:70036636..70039387 | Cyto | 3:02 |
| Sobic.001G419700 | SbHSP70-10 | Chr001:70042031..70044958 | Cyto | 2:01 |
| Sobic.001G419801 | SbHSP70-11 | Chr001:70045891..70048830 | Cyto | 2:01 |
| Sobic.001G419900 | SbHSP70-12 | Chr001:70053441..70055986 | Cyto | 2:01 |
| Sobic.001G420100 | SbHSP70-13 | Chr001:70059899..70062529 | Cyto | 2:01 |
| Sobic.001G454000 | SbHSP70-14 | Chr001:73039652..73043648 | Mito | 2:01 |
| Sobic.002G008000 | SbHSP70-15 | Chr002:746212..747519 | ER | 1:00 |
| Sobic.002G249800 | SbHSP70-16 | Chr002:63685205..63688420 | Mito | 5:04 |
| Sobic.003G178101 | SbHSP70-17 | Chr003:45559241..45561537 | Nucl | 5:04 |
| Sobic.003G350700 | SbHSP70-18 | Chr003:67021779..67024939 | Cyto | 2:01 |
| Sobic.003G378700 | SbHSP70-19 | Chr003:69252194..69254407 | ER | 1:00 |
| Sobic.004G011700 | SbHSP70-20 | Chr004:962795..966975 | ER | 9:08 |
| Sobic.004G263500 | SbHSP70-21 | Chr004:60838256..60844705 | Mito | 13:12 |
| Sobic.006G055600 | SbHSP70-22 | Chr006:39888060..39890944 | Cyto | 4:03 |
| Sobic.008G038700 | SbHSP70-23 | Chr008:3664254..3666130 | Cyto | 2:01 |
| Sobic.008G088200 | SbHSP70-24 | Chr008:18774401..18779411 | Chlo | 8:07 |
| Sobic.008G136000 | SbHSP70-25 | Chr008:56470318..56474560 | Cyto | 2:01 |
| Sobic.009G066900 | SbHSP70-26 | Chr009:7308207..7315031 | Cyto | 9:08 |
| Sobic.009G067000 | SbHSP70-27 | Chr009:7341758..7347966 | Cyto | 9:08 |
| Sobic.009G163900 | SbHSP70-28 | Chr009:52032825..52036139 | Cyto | 2:01 |
| Sobic.009G254800 | SbHSP70-29 | Chr009:58879456..58881262 | Cyto | 3:02 |
| Sobic.010G230600 | SbHSP70-30 | Chr010:57307086..57313310 | Nucl | 9:08 |

Cyto (cytoplasmic) Chlo(Chloroplasmic), Nucl(Nuclear),Mito(mitochondrial), ER( Endoplasmic Reticulum) Lp (localization predicted), Chro(Chromosome number), CDS( coding sequence), MW (Molecular Weight in Kilo Daltons), A.A (number of amino acids), pI (Isoelectric point), AI (Aliphatic Index), II (Instability Index), and GRAVY (Grand Average of hydropathicity Index)].

The intronless SbHSP70 genes share 6.66% of SbHSP70, while those with introns share 93.34%. The majority of SbHSP70s protein localized in the cytoplasm (17) and contains 1-8 introns, however, other SbHSP70 proteins localized in other organelles vary from 0 or 12 introns (Table-4). Similar findings were reported in different plants with varying intron numbers. For instance, in soybean (0-13) [45], *Arabidopsis thaliana, Phaseolus vulgaris*, and Popular HSP70 genes [34], some intronless genes were identified in the different TaHSP subfamilies in *Triticum aestivum* [17].

The structure of a gene determines its coding potential and can also give hints about the ancestry of genes since genes with similar structures probably evolved from a common ancestor [18]. The effect of introns on the transcription of genes is an evolutionarily conserved feature, being exhibited by such diverse organisms as yeast, plants, and mammals. Intron-containing genes are often transcribed more efficiently than non-intronic genes, and therefore, the presence of introns in a gene is generally associated with an increase in protein production mediated through many different mechanisms ranging from the increase of transcription and translation rates to the improvement of mRNA stability and folding [24].

### 3.3. Protein Motif and Cis Regulatory Element Prediction of HSP70 Genes

MEME was used to analyze the conserved motifs of different groups of protein sequences, and 10 conserved motifs were obtained (Figure-2), which were named Motifs 1-10 (Table-2).

**Figure 2:**
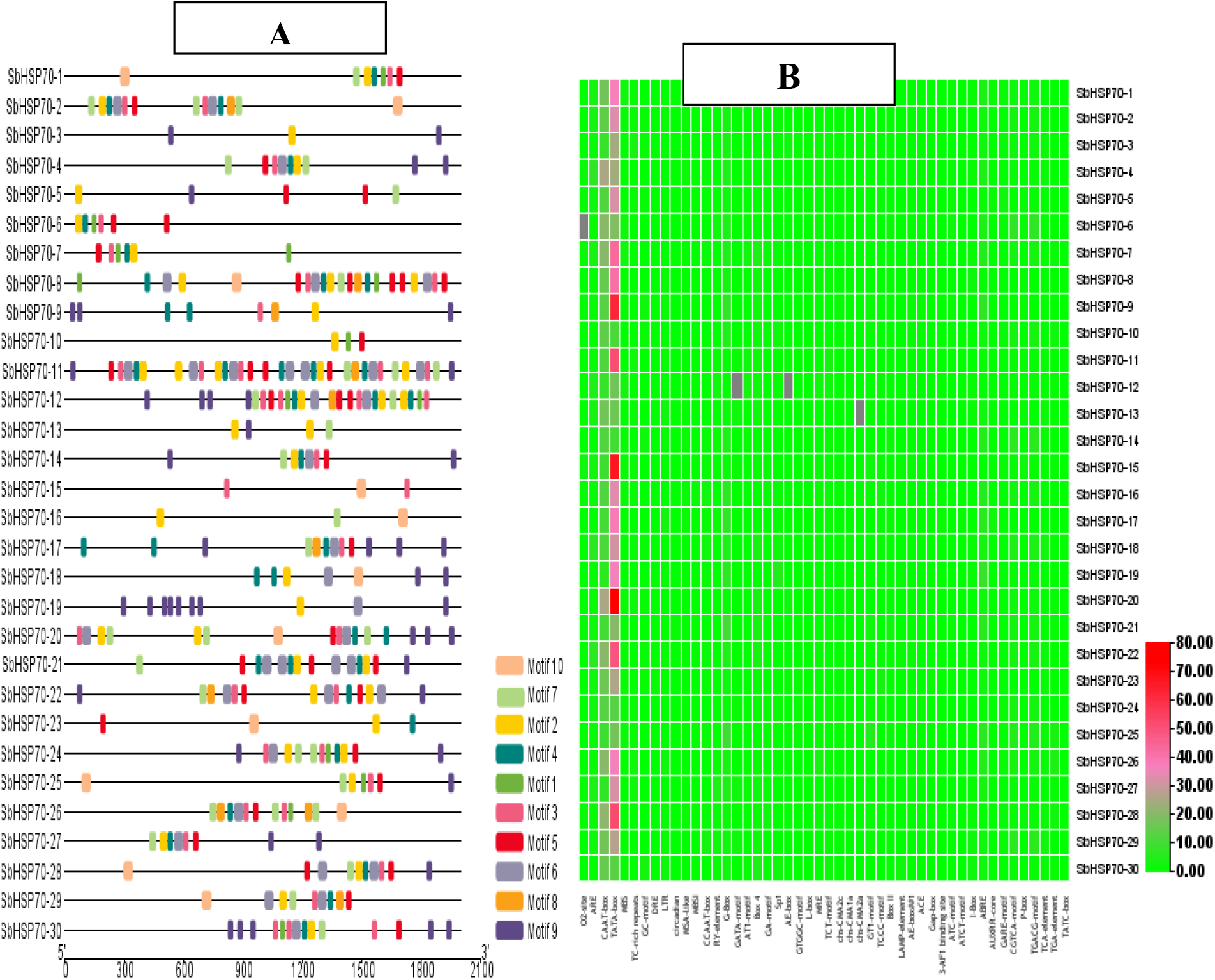
The Schematic representation of the identified motifs and Cis-regulatory elements of HSP70 gene in sorghum.

In the promoter region analysis, upstream of the transcription start site of the HSP70 gene family contained a total of 2,219 different cis-acting elements which are categorized as growth hormone-related elements (286) light-responsive elements (335), promoter-related elements (1,411), development/cell cycle-related elements (7), drought-related elements(20), metabolism-related elements(72) seed-specific regulation related elements(16) binding site related elements (18), temperature-related elements(19), stress defense-related elements(20) and anoxic related elements (15) were identified as computed in PLANT CARE enriched cis-acting element analysis database (Table-6). The detail of cis-acting elements is presented in Figure-2B.

Among the cis-acting elements discovered, the promoter enhancing elements (1411) are the leading and followed by light-responsive elements (335). All SbHSP70 genes contained growth hormone-related elements, promoter enhancer-related elements, and light-responsive related elements, at least two and other cis-acting elements vary among those genes. Average number of cis-acting elements per SbHSP70 gene was 73.97% and the highest number (117) of cis-elements was found on SbHSP70-20 followed by SbHSP70-9 (103) SbHSP70-15 (102) SbHSP70-22 (101) while the least (48) was on SbHSP70-24 (Table-6). The current results are in line with those of the previous report of HSP70 gene in Glycine *max* [45], PvHSP70 gene in *Phaseolus vulgaris*.

### 3.4. Physico-Chemical Properties of HSP70 Protein in Sorghum bicolor

The analyzed physical and chemical properties of HSP70 protein in sorghum are presented in Table-3. The functional diversity of HSP70 isoforms could be realized from the wide scope of their MW (33.01 kDa to 103.79 kDa) and the total number of amino acids in various HSP70 proteins that went from 299 amino acid length (SbHSP70-11) to 928 amino acids length in (SbHSP70-21). The analysis of the current study uncovered that all HSP70 protein families with record ID (SbHSP70-14, SbHSP70-17, SbHSP70-21, SbHSP70-26, SbHSP70-27, and SbHSP70-30) are unstable as determined by instability index result. The isoelectric point indicates that among the 30 HSP70 protein families recognized, 23 of the families have an acidic nature while the four families are basic and the rests are neutral. The aliphatic index for all HSP70 families is greater than 77.25 and therefore all identified HSP70 protein families have thermo stability. Whereas, lower GRAVY values of HSP70 indicate its hydrophilic nature.

In terms of Physicochemical characteristics such as gene size, protein length, and molecular weight (kDa) of the genes HSP70-1, HSP70-2, HSP70-3, HSP70-4, HSP70-13, HSP70-14, HSP70-16, HSP70-17, HSP70-18, HSP70-19, HSP70-20, HSP70-24, HSP70-25, and HSP70-28 were very similar to each other (Table-3). Similarly, HSP70-5, HSP70-6, HSP70-7, HSP70-8, HSP70-9, HSP70-10, HSP70-12, and HSP70-22 genes showed very similar physicochemical characteristics in terms of gene size, protein length, molecular weight (kDa), and PI (Table-3). Hence, it could be theorized that the molecular variation inside the characterized genes will have a fundamental role in biochemical and physiological functions that gives competitor gene-based markers, which show a nearby relationship with the trait of interest.

### 3.5. Predicted Proteins Sub-Cellular Localization

The analysis undertaken by cello life for subcellular localization of HSP70s in sorghum uncovered that these proteins were dispersed into five areas, for example, cytoplasmic (Cyto), endoplasmic reticulum (ER), chloroplast (Chlo), nuclear (Nucl), and mitochondrial (Mito). Most extreme proteins were found in the cytoplasmic/cytosolic (17), followed by four in chloroplast and mitochondrial. The endoplasmic reticulum represents three and the nucleus offers two genes (Table-3). In the current investigation of *Sorghum bicolor*, an aggregate of 17 cytosolic HSP70s proteins were discovered, which is similarly higher than the previous report in *Oryza sativa* [12] and *Arabidopsis thaliana* five [6] and lower than as contrasted in *Glycine max*, that is, 34 [34, 35, 46].

### 3.6. Gene Localization and Gene Duplication Analysis

The distinguished 30 HSP70 genes were conveyed on eight chromosomes of Sorghum bicolor (Figure 3). A large portion of the HSP70 genes were available on chromosome one (14 genes) and chromosome nine (four genes), while every one of the leftover chromosomes had one or three genes.

**Figure 3:**
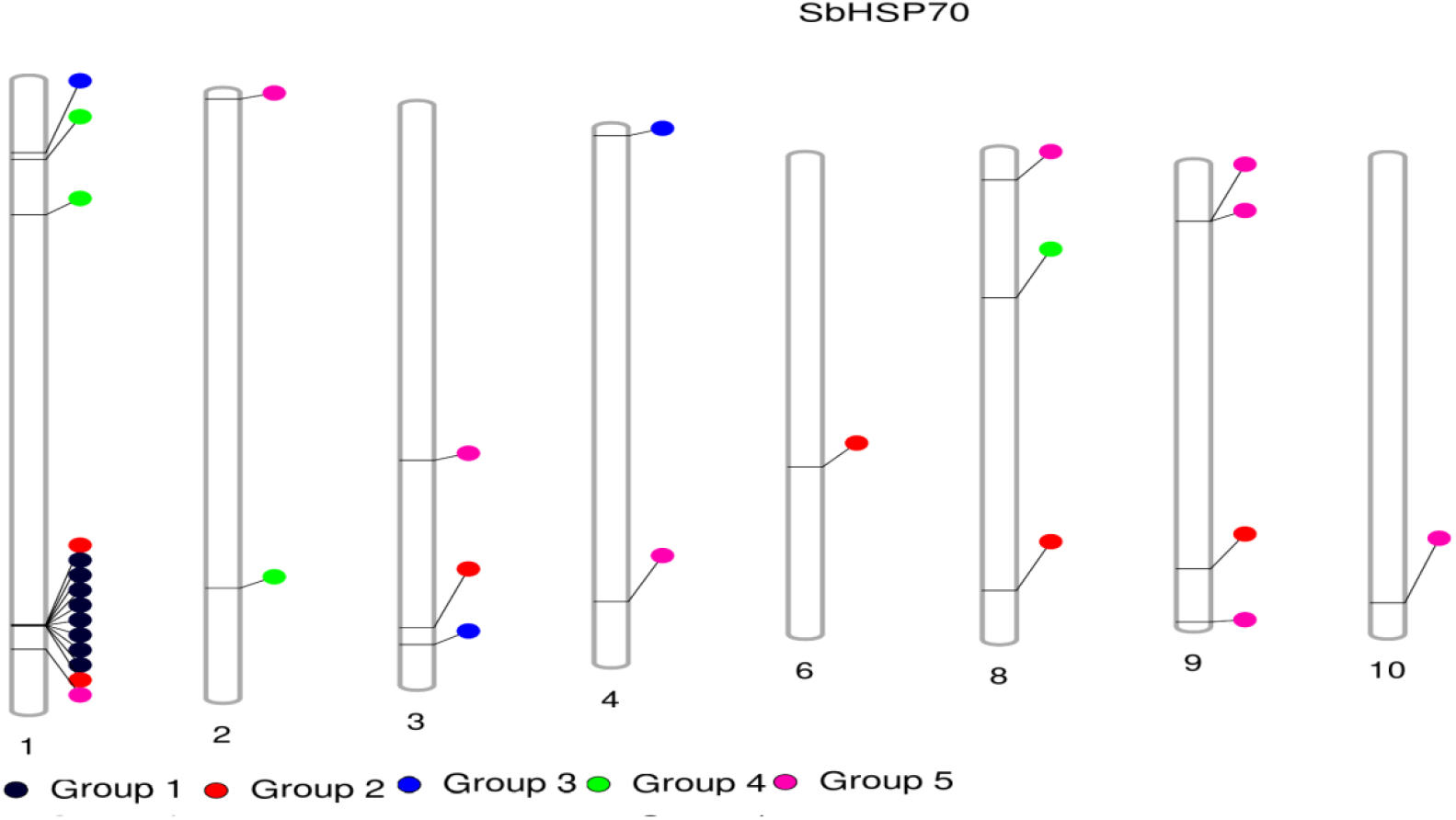
Distribution of HSP70 Genes on Sorghum bicolor Chromosome.

Both pair and segmental duplications add to the creation of gene families during advancement. In this way, potential duplication occasions of HSP70 gene were analized by insilico. Moreover, none of the genes were recommended to be the results of segmental duplication. In light of the outcomes, it could be inferred that pair duplication assumed a significant part in the extension of the HSP70 family in Sorghum bicolor.

To examine the molecular transformative pace of copied gene combines, then the nonsynonymous substitution (Ka) and synonymous substitution (Ks) proportions were determined utilizing Ka/Ks calculator in TBtool. In light of a pace of 6.1 x 10^-9^ replacements for each site each year, the difference time (T) was determined as T= Ks/(2 x 6.1 10^-9^) x 10^-6^ million years prior (Mya) (Table 5).

**Table 5:** Synonymous and Non-Synonymous Substitution Rates.

| Paralogs Gene Pairs |  | Ka | Ks | Ka/Ks | Time(MYA) |
| --- | --- | --- | --- | --- | --- |
| Sobic.003G350700 | Sobic.006G055600 | 0.014914382 | 0.372338146 | 0.040056013 | 0.0002 |
| Sobic.001G420100 | Sobic.008G136000 | 0.048386205 | 0.693205314 | 0.069800684 | 0.0003 |
| Sobic.001G419500 | Sobic.001G419600 | 0.105404002 | 0.468741704 | 0.22486585 | 0.0002 |
| Sobic.001G419000 | Sobic.001G419801 | 0.092001205 | 0.272097516 | 0.338118504 | 0.0001 |
| Sobic.003G378700 | Sobic.004G011700 | 0.202893273 | 1.076531965 | 0.188469344 | 0.0005 |
| Sobic.001G129000 | Sobic.008G088200 | 0.091415608 | 1.319369683 | 0.069287334 | 0.0007 |
| Sobic.001G193500 | Sobic.002G249800 | 0.15131001 | 2.809353502 | 0.05385937 | 0.0014 |
| Sobic.002G008000 | Sobic.008G038700 | 0.411522184 | 0.742245829 | 0.554428423 | 0.0004 |
| Sobic.009G066900 | Sobic.009G067000 | 0.047744901 | 0.488664962 | 0.097704777 | 0.0002 |

**Table 6:** Cis-acting elements analysis results as computed in PLANTCARE.

| Gene Name | GH R | PER | LRR | MBR | SDR | CCR | BSR | DRR | TR | SSR | ANR | Total | % |
| --- | --- | --- | --- | --- | --- | --- | --- | --- | --- | --- | --- | --- | --- |
| SbHSP70-1 | 10 | 54 | 12 | 3 | 1 | 0 | 0 | 2 | 0 | 0 | 0 | 82 | 7.45 |
| SbHSP70-2 | 9 | 47 | 4 | 3 | 1 | 1 | 0 | 1 | 1 | 0 | 1 | 68 | 6.18 |
| SbHSP70-3 | 8 | 32 | 11 | 1 | 3 | 0 | 0 | 0 | 0 | 0 | 1 | 56 | 5.09 |
| SbHSP70-4 | 7 | 49 | 11 | 8 | 0 | 1 | 1 | 0 | 0 | 0 | 0 | 77 | 7.00 |
| SbHSP70-5 | 7 | 44 | 4 | 3 | 1 | 0 | 0 | 0 | 0 | 0 | 1 | 60 | 5.45 |
| SbHSP70-6 | 20 | 39 | 9 | 1 | 0 | 0 | 0 | 2 | 0 | 2 | 0 | 73 | 6.64 |
| SbHSP70-7 | 12 | 63 | 15 | 4 | 0 | 0 | 1 | 2 | 0 | 0 | 0 | 97 | 8.82 |
| SbHSP70-8 | 2 | 56 | 9 | 0 | 0 | 0 | 0 | 0 | 1 | 0 | 0 | 68 | 6.18 |
| SbHSP70-9 | 8 | 75 | 14 | 2 | 0 | 0 | 0 | 1 | 1 | 0 | 2 | 103 | 9.36 |
| SbHSP70-10 | 11 | 25 | 13 | 0 | 0 | 0 | 0 | 0 | 0 | 0 | 1 | 50 | 4.55 |
| SbHSP70-11 | 5 | 63 | 9 | 1 | 2 | 0 | 0 | 0 | 1 | 1 | 0 | 82 | 7.45 |
| SbHSP70-12 | 3 | 25 | 12 | 4 | 0 | 0 | 0 | 3 | 0 | 2 | 0 | 49 | 4.45 |
| SbHSP70-13 | 4 | 33 | 7 | 3 | 1 | 0 | 0 | 0 | 2 | 0 | 0 | 50 | 4.55 |
| SbHSP70-14 | 18 | 22 | 7 | 0 | 0 | 1 | 1 | 1 | 1 | 1 | 2 | 54 | 4.91 |
| SbHSP70-15 | 6 | 76 | 12 | 5 | 1 | 0 | 1 | 1 | 0 | 0 | 0 | 102 | 9.27 |
| SbHSP70-16 | 11 | 45 | 15 | 2 | 1 | 1 | 3 | 1 | 0 | 0 | 0 | 79 | 7.18 |
| SbHSP70-17 | 8 | 48 | 13 | 0 | 0 | 1 | 1 | 0 | 3 | 1 | 0 | 75 | 6.82 |
| SbHSP70-18 | 14 | 42 | 13 | 6 | 1 | 0 | 2 | 0 | 0 | 0 | 1 | 79 | 7.18 |
| SbHSP70-19 | 10 | 42 | 13 | 1 | 0 | 0 | 1 | 0 | 1 | 0 | 0 | 68 | 6.18 |
| SbHSP70-20 | 7 | 98 | 7 | 1 | 2 | 0 | 0 | 1 | 0 | 0 | 1 | 117 | 10.64 |
| SbHSP70-21 | 13 | 30 | 19 | 3 | 0 | 1 | 0 | 0 | 0 | 0 | 0 | 66 | 6.00 |
| SbHSP70-22 | 9 | 69 | 13 | 6 | 0 | 0 | 1 | 1 | 1 | 0 | 1 | 101 | 9.18 |
| SbHSP70-23 | 3 | 42 | 9 | 2 | 0 | 0 | 1 | 1 | 1 | 0 | 1 | 60 | 5.45 |
| SbHSP70-24 | 8 | 26 | 10 | 2 | 1 | 0 | 0 | 1 | 0 | 0 | 0 | 48 | 4.36 |
| SbHSP70-25 | 20 | 26 | 18 | 1 | 1 | 0 | 1 | 1 | 0 | 0 | 0 | 68 | 6.18 |
| SbHSP70-26 | 17 | 56 | 8 | 1 | 2 | 0 | 0 | 0 | 2 | 1 | 0 | 87 | 7.91 |
| SbHSP70-27 | 6 | 39 | 11 | 5 | 2 | 0 | 0 | 0 | 0 | 4 | 0 | 67 | 6.09 |
| SbHSP70-28 | 7 | 73 | 9 | 2 | 0 | 1 | 1 | 0 | 1 | 1 | 0 | 95 | 8.64 |
| SbHSP70-29 | 17 | 44 | 11 | 2 | 0 | 0 | 3 | 0 | 1 | 2 | 0 | 80 | 7.27 |
| SbHSP70-30 | 6 | 28 | 17 | 0 | 0 | 0 | 0 | 1 | 2 | 1 | 3 | 58 | 5.27 |
| Total | 286 | 1411 | 335 | 72 | 20 | 7 | 18 | 20 | 19 | 16 | 15 | 2219 | 73.97 |

GHR(Growth Hormone Related), PER( Promoter Enhancer Related), LRR (Low-Temperature Related), MBR (Metabolism Related), SDR( Stress Defense Related), CCR( Cell Cycle Related), BSR( Binding Site Related), DRR (Drought Response Related), TR ( Temperature Related), SSR(Seed Specific Regulation), ANR (Anoxic Related)

Gene duplications assume a significant part in advancement as duplications cause genes to create gene families [15]. Truth be told, it has been recommended that couple and segmental duplications have been the essential driving the wellspring of evolution as these occasions lead to extension of gene families, and age of proteins with novel capacities [5]. Tandem duplication includes the duplication of at least two genes situated on a similar chromosome, while segmental duplication alludes to the marvel when genes having a place in a similar clade, however, situated on various chromosomes are duplicated[20].

In the current study, an aggregate of eighteen (18/30; 60 %) Sorghum bicolor HSP70 genes were demonstrated to be copied (Table 5). Furthermore, the seven sets of a gene had all earmarks of being tandemly duplicated, which was perceived on chromosome number one (Figure 3). The remainder of the copied genes were all segmentally copied/duplicated.

The proportion of Ka and Ks replacement rate is a powerful strategy to research the specific imperative among copied gene sets [32]. Henceforth, in the current review, Ka, Ks, and Ka/Ks esteems for each pair of paralogous genes were determined (Table 4). On an essential level, the worth of Ka/Ks < 1 connotes the decontaminating option (negative option), Ka/Ks > 1 implies positive determination/selection, and Ka/Ks = 1 the system impartial option (20). At this point, 18 HSP70 genes were demonstrated to be copied. The Ka/Ks proportion for copied HSP70 genes went from 0.040056013 to 0.554428423. All HSP70 genes in the current analysis have Ka/Ks value < 1(Table 5).

### 3.7. Phylogenetic Analysis

Phylogeny helps in understanding how genes diverge during evolution. Therefore, in the current study, the phylogenetic analysis was performed to investigate the phylogenetic relationships of HSP70 protein families with homolog sequences from different plant species. The tree was constructed utilizing amino acid sequences from *Sorghum bicolor*, *Zea mays*, *Oryza sativa*, *Triticum aestivum*, *Saccharum officinarum*, *Panicum miliaceum*, and the model plant *Arabidopsis thaliana* with a total of 53 HSP70 protein families; following the Neighbor-Joining method. The bootstrap consensus tree deduced from 1000 replicates was taken to address the evolutionary history of the taxa investigated. Each HSP70 family gene were clustered together into eight groups (Figure 4). In general, phylogenetic relationships revealed that the sorghum HSP70 gene family is present in all groups and this presence ties its relationship closely to *Arabidopsis thaliana*, which is present in all groups except groups five and seven and most of the HSP70 proteins are far apart from HSP70 protein family in sorghum. In the phylogenetic tree, all members of HSP70 of *Sorghum bicolor* are well separated into distinct systems and they are grouped with proteins of other species, with a strong bootstrap values (Figure 4).

**Figure 4:**
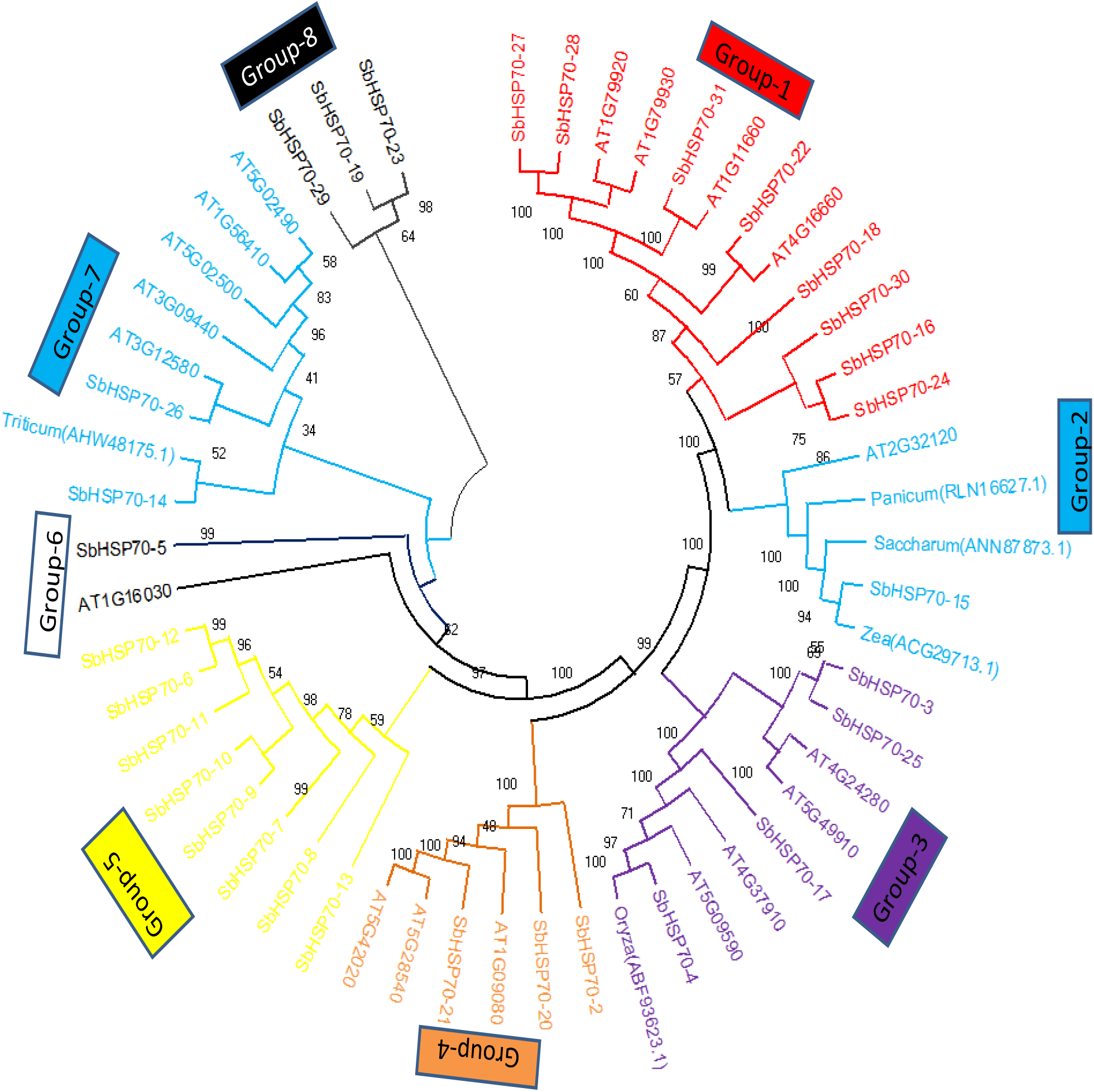
Phylogenetic tree (neighbor-joining) of the heat shock protein 70 from sorghum and other species.

## 4. CONCLUSIONS

With the increasing concerns about global warming and rising earth temperatures, it is essential to know about proteins that provide heat/stress tolerance in crop plants. The present study identified and characterized the HSP70 family in *Sorghum bicolor* genome. The Genome-wide assortment of *Sorghum bicolor* genomes for the identification of SbHSP70 revealed the presence of 30 genes. The different analyses performed disclosed their structural organization, subcellular localization, physicochemical properties, cis-acting elements, phylogenetic, and understress conditions. This study provides further information for the functional characterization of HSP70 and helps to understand the mechanisms of abiotic stress tolerance under diverse stress conditions in *Sorghum bicolor*.

## Acknowledgements

We are grateful to Department of Applied Biology, School of Applied Natural Science, Adama Science and Technology University for kindly providing us with an accessible internate aceess and Mekdela Amba University for providing the chance to carry out the research for the corresponding auther.

## Authors’ contribution

The corresponding author of this manuscript has written the whole document as well as conducted the analysis in silico and the co-author Mulugeta Kebede reviewed the document and guided the author during the whole work.

## Availability of Data and Materials

The data sets used or analyzed during the current study for sorghum genes are available at Phytozome V3.1.1 (Cereal grass) database(https://phytozome-next.jgi.doe.gov/info/Sbicolor_v3_1_1), with phytozome accession number of Sobic.001G118600, Sobic.001G129000, Sobic.001G193500, Sobic.001G418600, Sobic.001G419000, Sobic.001G419300, Sobic.001G419400, Sobic.001G419500, Sobic.001G419600, Sobic.001G419700, Sobic.001G419801, Sobic.001G419900, Sobic.001G420100, Sobic.001G454000, Sobic.002G008000, Sobic.002G249800, Sobic.003G178101, Sobic.003G350700, Sobic.003G378700, Sobic.004G011700, Sobic.004G263500, Sobic.006G055600, Sobic.008G038700, Sobic.008G088200, Sobic.008G136000, Sobic.009G066900, Sobic.009G067000, Sobic.009G163900, Sobic.009G254800 and Sobic.010G230600 respectively. Inaddition, *Arabdopsis thaliana* (AT1G09080, AT1G11660, AT1G16030, AT1G56410, AT1G79920, AT1G79930, AT2G32120, AT3G09440, AT3G12580, AT4G16660, AT4G24280, AT4G37910, AT5G02490, AT5G02500, AT5G09590, AT5G28540, AT5G42020, AT5G49910), gene accession were also brosed from phytozome. Where as, *Triticum aestivum, Zea mays, Panicum miliaceum, Saccharum officinarum, and Oryza sativa* from NCBI with accession number of AHW48175.1, ACG29713.1, RLN16627.1, (ANN87873.1, ABF93623.1 repectively.

## Declaration

Ethics approval and consent to participate

Not applicable.

## Consent for Publication

Not applicable

## REFERENCES

1. Amelework BA, Shimelis HA, Laing MD, Ayele DG, Tongoona P, Mengistu F. Sorghum production systems and constraints and coping strategies under drought-prone agroecologies of Ethiopia. South African Journal of Plant and Soil. 2016, Jan 1;33(3):207–17.

2. Ananda G, Gleadow R, Norton S, Furtado A, Henry R. Determination of Phylogenetic Relationships of the Genus Sorghum Using Nuclear and Chloroplast Genome Assembly. Multidisciplinary Digital Publishing Institute Proceedings. 2019;36(1):17.

3. Bailey TL, Boden M, Buske FA, Frith M, Grant CE, Clementi L, Ren J, Li WW, Noble WS. MEME SUITE: tools for motif discovery and searching. Nucleic acids research. 2009 Jul 1;37(suppl_2):W202–8.

4. Boone AN, Vijayan, MM. Constitutive heat shock protein 70 (HSC70) expression in rainbow trout hepatocytes: effect of heat shock and heavy metal exposure. Comparative Biochemistry and Physiology Part C: Toxicology & Pharmacology. 2002 Jun 1;132(2):223–33.

5. Cannon SB, Mitra A, Baumgarten A, Young ND, May G. The roles of segmental and tandem gene duplication in the evolution of large gene families in Arabidopsis thaliana. BMC plant biology. 2004 Dec;4(1):1–21.

6. Chaudhary R, Baranwal VK, Kumar R, Sircar D, Chauhan H. Genome-wide identification and expression analysis of Hsp70, Hsp90, and Hsp100 heat shock protein genes in barley under stress conditions and reproductive development. Functional & integrative genomics. 2019 Nov;19(6):1007–22.

7. Duan YH, Guo J, Ding K, Wang SJ, Zhang H, Dai XW, Chen YY, Govers F, Huang LL, Kang ZS. Characterization of a wheat HSP70 gene and its expression in response to stripe rust infection and abiotic stresses. Molecular biology reports. 2011 Jan;38(1):301–7.

8. Guo M, Liu JH, Ma X, Zhai YF, Gong ZH, Lu MH. Genome-wide analysis of the Hsp70 family genes in pepper (Capsicum annuum L.) and functional identification of CaHsp70-2 involvement in heat stress. Plant Science. 2016 Nov 1;252:246–56.

9. Guo M, Zhai YF, Lu JP, Chai L, Chai WG, Gong ZH, Lu MH. Characterization of CaHsp70-1, a pepper heat-shock protein gene in response to heat stress and some regulation exogenous substances in Capsicum annuum L. International journal of molecular sciences. 2014 Nov;15(11):19741–59.

10. Gupta SC, Sharma A, Mishra M, Mishra RK, Chowdhuri DK. Heat shock proteins in toxicology: how close and how far?. Life sciences. 2010 Mar 13;86(11-12):377–84.

11. Guy CL, Li QB. The organization and evolution of the spinach stress 70 molecular chaperone gene family. The Plant Cell. 1998 Apr;10(4):539–56.

12. Haag J. Molecular and biochemical enhancement of chlorophyll in sports turf. Lulu. com; 2019 Feb 21.

13. Hirayama T, Shinozaki K. Research on plant abiotic stress responses in the post-genome era: Past, present and future. The Plant Journal. 2010 Mar;61(6):1041–52.

14. Hu B, Jin J, Guo AY, Zhang H, Luo J, Gao G. GSDS 2.0: an upgraded gene feature visualization server. Bioinformatics. 2015 Apr 15;31(8):1296–7.

15. Hu W, Hu G, Han B. Genome-wide survey and expression profiling of heat shock proteins and heat shock factors revealed overlapped and stress specific response under abiotic stresses in rice. Plant Science. 2009 Apr 1;176(4):583–90.

16. Jiang M, Chu Z. Comparative analysis of plant MKK gene family reveals novel expansion mechanism of the members and sheds new light on functional conservation. Bmc Genomics. 2018 Dec;19(1):1–8.

17. Krishnamurthy L, Serraj R, Hash CT, Dakheel AJ, Reddy BV. Screening sorghum genotypes for salinity tolerant biomass production. Euphytica. 2007 Jul;156(1):15–24.

18. Kumar A, Sharma S, Chunduri V, Kaur A, Kaur S, Malhotra N, Kumar A, Kapoor P, Kumari A, Kaur J, Sonah H. Genome-wide Identification and Characterization of Heat Shock Protein Family Reveals Role in Development and Stress Conditions in Triticum aestivum L. Scientific reports. 2020 May 12;10(1):1–2.

19. Liu J, Pang X, Cheng Y, Yin Y, Zhang Q, Su W, Hu B, Guo Q, Ha S, Zhang J, Wan H. The Hsp70 gene family in Solanum tuberosum: genome-wide identification, phylogeny, and expression patterns. Scientific reports. 2018 Nov 9;8(1):1–1.

20. Liu Y, Jiang H, Chen W, Qian Y, Ma Q, Cheng B, Zhu S. Genome-wide analysis of the auxin response factor (ARF) gene family in maize (Zea mays). Plant Growth Regulation. 2011 Apr;63(3):225–34.

21. Lynch M, Conery JS. The evolutionary fate and consequences of duplicate genes. science. 2000 Nov 10;290(5494):1151–5.

22. Maimbo M, Ohnishi K, Hikichi Y, Yoshioka H, Kiba A. Induction of a small heat shock protein and its functional roles in Nicotiana plants in the defense response against Ralstonia solanacearum. Plant physiology. 2007 Dec;145(4):1588–99.

23. Marchler-Bauer A, Bo Y, Han L, He J, Lanczycki CJ, Lu S, Chitsaz F, Derbyshire MK, Geer RC, Gonzales NR, Gwadz M. CDD/SPARCLE: functional classification of proteins via subfamily domain architectures. Nucleic acids research. 2017 Jan 4;45(D1):D200–3.

24. Moabbi AM, Agarwal N, El Kaderi B, Ansari A. Role for gene looping in intron-mediated enhancement of transcription. Proceedings of the National Academy of Sciences. 2012 May 29;109(22):8505–10.

25. Mosa KA, Ismail A, Helmy M. Introduction to plant stresses. InPlant stress tolerance 2017 (pp. 1–19). Springer, Cham.

26. Mulaudzi-Masuku T, Mutepe RD, Mukhoro OC, Faro A, Ndimba B. Identification and characterization of a heat-inducible Hsp70 gene from S orghum bicolor which confers tolerance to thermal stress. Cell Stress and Chaperones. 2015 Sep;20(5):793–804.

27. Nagaraju M, Reddy PS, Kumar SA, Kumar A, Rajasheker G, Rao DM, Kishor PK. Genome-wide identification and transcriptional profiling of small heat shock protein gene family under diverse abiotic stress conditions in Sorghum bicolor (L.). International journal of biological macromolecules. 2020 Jan 1;142:822–34.

28. Ndimba BK, Thomas LA, Ngara R. Sorghum 2-dimensional proteome profiles and analysis of Hsp70 expression under salinity stress. Agriculture and Natural Resources. 2010 Oct 30;44(5):768–75.

29. Ngara R, Ndimba R, Borch-Jensen J, Jensen ON, Ndimba B. Identification and profiling of salinity stress-responsive proteins in Sorghum bicolor seedlings. Journal of Proteomics. 2012 Jul 16;75(13):4139–50.

30. Ohama N, Sato H, Shinozaki K, Yamaguchi-Shinozaki K. Transcriptional regulatory network of plant heat stress response. Trends in plant science. 2017 Jan 1;22(1):53–65.

31. Ré MD, Gonzalez C, Escobar MR, Sossi ML, Valle EM, Boggio SB. Small heat shock proteins and the postharvest chilling tolerance of tomato fruit. Physiologia plantarum. 2017 Feb;159(2):148–60.

32. Rehman S, Jørgensen B, Aziz E, Batool R, Naseer S, Rasmussen SK. Genome wide identification and comparative analysis of the serpin gene family in brachypodium and barley. Plants. 2020 Nov;9(11):1439.

33. Rodziewicz P, Swarcewicz B, Chmielewska K, Wojakowska A, Stobiecki M. Influence of abiotic stresses on plant proteome and metabolome changes. Acta Physiologiae Plantarum. 2014 Jan;36(1):1–9.

34. Sung DY, Vierling E, Guy CL. Comprehensive expression profile analysis of the Arabidopsis Hsp70 gene family. Plant physiology. 2001 Jun 1;126(2):789–800.

35. Sarkar NK, Kundnani P, Grover A. Functional analysis of Hsp70 superfamily proteins of rice (Oryza sativa). Cell stress and Chaperones. 2013 Jul;18(4):427–37.

36. Segui-Simarro JM, Testillano PS, Risueno MC. Hsp70 and Hsp90 change their expression and subcellular localization after microspore embryogenesis induction in Brassica napus L. Journal of structural biology. 2003 Jun 1;142(3):379–91.

37. Shukla V, Upadhyay RK, Tucker ML, Giovannoni JJ, Rudrabhatla SV, Mattoo AK. Transient regulation of three clustered tomato class-I small heat-shock chaperone genes by ethylene is mediated by SlMADS-RIN transcription factor. Scientific reports. 2017 Jul 25;7(1):1–2.

38. Taiz L, Zeiger E, Møller IM, Murphy A. Plant physiology and development. Sinauer Associates Incorporated; 2015.

39. Tari I, Laskay G, Takács Z, Poór P. Response of sorghum to abiotic stresses: A review. Journal of Agronomy and Crop Science. 2013 Aug;199(4):264–74.

40. Tyedmers J, Mogk A, Bukau B. Cellular strategies for controlling protein aggregation. Nature reviews Molecular cell biology. 2010 Nov;11(11):777–88.

41. Vacchina P, Norris-Mullins B, Carlson ES, Morales MA. A mitochondrial HSP70 (HSPA9B) is linked to miltefosine resistance and stress response in Leishmania donovani. Parasites & vectors. 2016 Dec;9(1):1–5.

42. Vega VL, Rodríguez-Silva M, Frey T, Gehrmann M, Diaz JC, Steinem C, Multhoff G, Arispe N, De Maio A. Hsp70 translocates into the plasma membrane after stress and is released into the extracellular environment in a membrane-associated form that activates macrophages. The Journal of Immunology. 2008 Mar 15;180(6):4299–307.

43. Wang W, Vinocur B, Shoseyov O, Altman A. Role of plant heat-shock proteins and molecular chaperones in the abiotic stress response. Trends in plant science. 2004 May 1;9(5):244–52.

44. Waters ER. The evolution, function, structure, and expression of the plant sHSPs. Journal of experimental botany. 2013 Jan 1;64(2):391–403.

45. Zhang J, Liu B, Li J, Zhang L, Wang Y, Zheng H, Lu M, Chen J. Hsf and Hsp gene families in Populus: genome-wide identification, organization and correlated expression during development and in stress responses. BMC genomics. 2015 Dec;16(1):1–9.

46. Zhang L, Zhao HK, Dong QL, Zhang YY, Wang YM, Li HY, Xing GJ, Li QY, Dong YS. Genome-wide analysis and expression profiling under heat and drought treatments of HSP70 gene family in soybean (Glycine max L.). Frontiers in plant science. 2015 Sep 25;6:773.

47. Zhu X, Zhao X, Burkholder WF, Gragerov A, Ogata CM, Gottesman ME, Hendrickson WA. Structural analysis of substrate binding by the molecular chaperone DnaK. Science. 1996 Jun 14;272(5268):1606–14.

